# Assessment of the bound conformation of Bombesin to the BB1 and BB2 Receptors

**DOI:** 10.1101/2023.07.18.549617

**Authors:** Guillem Vila-Julià, Jaime Rubio-Martinez, Juan J. Perez

## Abstract

Bombesin is an endogenous peptide involved in a wide spectrum of physiological activities ranging from satiety, control of circadian rhythm and thermoregulation in the central nervous system, to stimulation of gastrointestinal hormone release, activation of macrophages and effects on development in peripheral tissues. Actions of the peptide are mediated through the two high affinity G-protein coupled receptors BB1 and BB2. Under pathophysiological conditions, these receptors are overexpressed in many different types of tumors, such as prostate cancer, breast cancer, small and non-small cell lung cancer and pancreatic cancer. This knowledge has been used for designing cell markers, but it has not been yet exploited for therapeutical purposes. Despite the enormous biological interest of the peptide, little is known about the stereochemical features that contribute to their activity. On the one hand, mutagenesis studies identified a few receptor residues important for high bombesin affinity and on the other, a few studies focused on the relevance of diverse residues of the peptide for receptor activation. Models of the peptide bound to BB1 and BB2 can be helpful to improve our understanding of the stereochemical features granting bombesin activity. Accordingly, the present study describes the computational process followed to construct such models from models of the peptide and its receptors by means of Steered Molecular Dynamics. Present results provide new insights into the structure-activity relationships of bombesin and its receptors, as well as render an explanation for the differential binding affinity observed towards the BB1 and BB2 receptors. Finally, these models can be further exploited to help for designing novel small molecule peptidomimetics with improved pharmacokinetics profile.

**AUTHOR SUMMARY:** The goal of the present work is to construct models of bombesin bound to its receptors BB1 and BB2. The work represents an attempt to conceal experimental information available for bombesin activity on key residues of the sequence, as well as on specific residues in the receptors derived from site-directed mutagenesis with its structure. For this purpose, models of the two receptors were constructed homology using endothelin B as template and a model of bombesin structure in solution. Next, bombesin was docked onto each of the two receptors by means of Steered Molecular Dynamics, by pulling the peptide into the receptor using a constant force. Ten trials were performed on each receptor. After each trial, the resulting complex was relaxed using a 200 ns MD trajectory. In addition, the binding free energy was computed by means of the MMPBSA method for each of these simulations. Next, residue contributions to the binding free energy permitted to select the most suitable complex by comparison of their contributions to their importance deduced from experimental results. The best-fitted complex for each receptor was subject of a 2 μs MD simulation that permitted to compute a difference of the binding free energy of the peptide that agrees well with pharmacology data. Finally, a study of the binding free energy contributions per residue permitted to understand specific differences between the bound conformation of bombesin in two receptors that explain the observed differential affinity. Specifically, a non-conserved residue in ECL3 (Pro in BB1 and Thr in BB2) appears to be responsible of a differential interaction of Arg(6.58) with the peptide, in addition to provide an extra interaction with Gln7 of bombesin (in the case of Thr(ECL3)). These models permit to explain the differential pharmacological profile, despite the high sequence identity between the two receptors, shedding light into the structure-activity relationships of the peptide available.

## INTRODUCTION

Bombesin-like peptides comprise a large family of peptides characterized by the consensus sequence: Gly-X-Y-Met-NH_2_ (X=His or Ser; Y= Leu or Phe) at their C-terminus [1]. Despite these peptides were originally isolated from the skin of diverse amphibians, they are widely distributed in mammals accomplishing a wide spectrum of biological activities [2]. Thus, in the central nervous system, they are involved in satiety, control of circadian rhythm, thermoregulation and in peripheral tissues, in processes like the stimulation of gastrointestinal hormone release, activation of macrophages and effects on development [3, 4]. They are also involved in pathological processes, such as tumor growth and differentiation [5]. Their actions are mediated through three G-protein coupled receptors (GPCRs) known as BB1, BB2 and BB3 [6–8]. Under pathophysiological conditions these receptors are overexpressed in many different types of tumors, such as prostate cancer, breast cancer, small and non-small cell lung cancer and pancreatic cancer [9]. Despite the enormous biological interest of this family of peptides, little is known about the stereochemical features that contribute to their activity. Thus, a better understanding of the structure-activity relationships of the diverse members of the family is key for designing peptide analogues and peptidomimetics that can turn out to be novel therapeutic agents [10, 11].

Bombesin (Bn), a tetradecapeptide with the sequence: Glp-Gln-Arg-Leu-Gly-Asn-Gln-Trp-Ala-Val-Gly-His-Leu-Met-NH_2_ (Glp = pyroglutamic acid) is the most widely characterized member of the family. The peptide exhibits high affinity for the BB1 and BB2 receptors (K_i_=4 nM and 0.07 nM, respectively) [12], but does not bind to BB3. Synthesis and biological screening of diverse fragments and analogues permitted the identification of key residues involved in its activity. Thus, Bn(6-14) is the shortest fragment retaining full activity [12, 13] and residues Trp8, His12, Leu13 and Met14 were shown to be important for receptor binding [14, 15]. Moreover, Met14 is key for receptor activation, since its deletion transforms the peptide into a potent antagonist [16]. Furthermore, site directed mutagenesis provides additional information about key residues for binding and/or activity in the receptors. Thus, residues Gln123(3.32), Pro200(45.52) and Arg289(6.58) are known to be important for Bn binding to BB1(Residue numbering in parenthesis follows the Ballesteros–Weinstein notation system [22]). On the other hand, Gln120(3.32), Arg287(6.58) and Arg308(7.39) are known to be key for Bn binding to BB2 [17–19], along with residues Pro198(45.52), Ala307(7.38) and Phe184(ECL2) that were identified as binding modulators [18–21].

Regarding the conformational profile of the peptide, diverse spectroscopy and computational studies are available. The former include NMR [23–27], Infrared (IR) [28], Circular Dichroism (CD) and Fluorescence spectroscopy [29] in different solvents like water, dimethylsulfoxide (DMSO) or 2,2,2-trifluoroethanol-water mixtures. Specifically, early NMR reports in water and DMSO [23–25] describe the structure of Bn as a random coil. In contrast, NMR experiments in a 2,2,2-trifluoroethanol/water mixture (30% v/v) [26, 27] report that the segment Bn(6-14) displays a helical conformation, with residues 11 to 14 less sharply structured. Moreover, the same study also concludes that the first two N-terminal residues adopt an extended conformation, while the region between residues 3 and 5 exhibits a great deal of flexibility. IR [28], CD and Fluorescence studies [29] also confirm that the peptide adopts a helical conformation in lipid environments. On the other hand, computational studies show that the peptide attains a helical structure on the segment Bn(6-14) with a tendency to unwind at the C-term edge of the helix [30, 31]. Moreover, these studies also show a frequent occurrence of conformations that bring together the side chains of aromatic residues Trp8 and His12 closely, a fact that was considered important to explain the importance of these two residues for binding.

In order to improve our understanding on the stereochemical features granting Bn activity and to ascertain the reasons for the observed affinity difference to the BB1 and BB2 receptors, despite their high sequence identity in the orthosteric site, models of the peptide bound to each of the two receptors were produced. Such a models can help to understand the involvement of the diverse peptide residues either as secondary structure elements or as key actors in receptor recognition, providing information about key specific interactions [32]. Modeling involved the construction of the 3D structures of the two receptors by homology modeling that were used to dock bombesin onto them. Calculations were performed using a previously developed model of the peptide in solution [24,25] as starting conformation with its C-terminus facing the receptors orthosteric site. This assumption is funded on the experimental evidence outlined above, recognizing Bn(6-14) as the shortest fragment with activity and that the C-terminal residue Met14 as necessary for activation [4]. Further confirmation comes from the recently disclosed structures of the human BB2 receptor bound to the gastrin-releasing peptide and the synthetic agonist [D-Phe6, β-Ala11, Phe13, Nle14] Bn (6–14) [33]. In both structures, peptide ligands are bound with their C-terminus deeply located at the bottom of the orthosteric pocket.

Due to the uncertainties involving the induced fit that takes place on both, peptide and receptor upon binding, after considering alternative procedures [32], we resolved to carry out our study using Steered Molecular Dynamics (SMD) [34]. In this procedure, the peptide in a helical conformation is pulled with a constant force towards the receptor with its C-terminus facing the orthosteric site, allowing the peptide and the receptors to locally fit to each other. The outcomes of the present study are robust models of Bn-BB1 and Bn-BB2 complexes, together with an estimate of their respective binding free energy. Moreover, these models permit to understand the structural role of the diverse peptide residues in their interaction with the receptor and provide an explanation of the observed differential binding affinity of bombesin for the two receptors.

## RESULTS

Models of the BB1 and BB2 receptors were constructed by homology modelling using the structure of the human endothelin B receptor (PDB entry code 5X93) [32] as template. Endothelin B shares 32% identity with both, BB1 and BB2 receptors, and was chosen as template for being one of the closest GPCR whose 3D structure was available in the phylogenetic tree [58]. Sequence alignment of the BB1, BB2 receptors and the endothelin B receptor is shown in Figure S1 of the supplementary material and was used as guidance for the threading process. Sequence alignment of BB1 and BB2 shows that they exhibit a high identity in the orthosteric site region and much lower in the extracellular loops.

Rough models of the two receptors were obtained with MODELLER and subsequently refined using MD simulations, as explained in the methods section.^34^ Time evolution of the rmsd considering the Cα of the proteins for the three 500 ns MD replicas is shown in Figure S2 of the supplementary material. As can be seen, in both cases systems got equilibrated after 200 ns of the trajectory, as previously found [38]. Subsequent cluster analysis of the structures sampled during the MD process permitted to identify representative structures of the most populated cluster for each receptor that were considered the final receptor models, respectively. At the time this study was undertaken, no crystallographic structure of any of the bombesin receptors was available. However very recently, the structures of the human BB2 receptor bound to the gastrin-releasing peptide, the synthetic agonist [D-Phe6, β-Ala11, Phe13, Nle14] Bn (6– 14) or the antagonist PD176252 were released [33]. Comparison of the homology model of BB2 constructed in this work and the structure of BB2 with the antagonist PD176252 bound, solved by X-ray diffraction shows no significant differences in the orthosteric side. Actually, the RMSD in the TM(1-7) region is 1.40 Å. The largest differences are concentrated in N-terminus of TM1 and TM6, although differences may be attributed to an artifact of the crystallographic structure, due to the glycogen synthase inserted in the ICL3 used for crystallization purposes [33]. In contrast, when the extracellular loops are included in the comparison, the rmsd increases to 3.71 Å. This can be explained as the result of the large conformational fluctuations of the loops and to the removal of the N-terminus in our models to avoid artifacts during the sampling process.

Structures resulted from the different SMD trials correspond to bound conformations of bombesin to the respective receptors. Each of these structures were subsequently relaxed by means of 200 ns MD calculations. Structural stability of the complexes was monitored through the time evolution of the rmsd using Cα of the residues in the transmembrane region, referred to the starting structure and shown in Figures S3 and S4 of the supplementary material. Inspection of the Figures suggests that after a short stabilization time, no important structural reorganization occurs and all the complexes remain stable during the simulation time. Figures S5 and S6 of the supplementary material show the time evolution of ΔG_bind_ for each of the two receptors and final values are listed in Table 1, computed as the average values of the last 20ns of each replica. Most of the structures obtained at the end of the relaxation process subsequent to the SMD calculations show the Bn C-terminus sitting on the orthosteric site, partially unfolded as a consequence of the ligand-receptor induced fitting during the relaxation process and with the central segment in its initial helical conformation. The same features are also observed in the crystallographic structures of endothelin bound to the endothelin B receptor, as well as in the recently BB2-gastrin-releasing peptide complex recently disclosed, where the central peptide segment adopts a helical conformation and the C-terminus is partially unfolded [58].

**Table 1.** Binding free energy for the complexes bombesin-BB1 and bombesin-BB2 computed by means of the MMPBSA method, using the last 20 ns of the 200 ns relaxation MD trajectories (D01-D10) for each complex.

| Trajectory | AVERAGE $\Delta G_{\text{bind}}$ last 20ns from 200ns (kcal/mol) | |
| --- | --- | --- |
|  | Bn-BB1 | Bn-BB2 |
| D01 | -66.9 ± 0.3 | -105.1 ± 0.3 |
| D02 | -80.1 ± 0.3 | -85.5 ± 0.3 |
| D03 | -96.9 ± 0.4 | -94.7 ± 0.2 |
| D04 | -80.2 ± 0.2 | -73.4 ± 0.2 |
| D05 | -79.1 ± 0.2 | -69.3 ± 0.3 |
| D06 | -82.5 ± 0.3 | -92.1 ± 0.4 |
| D07 | -57.8 ± 0.4 | -102.2 ± 0.2 |
| D08 | -83.1 ± 0.3 | -77.5 ± 0.2 |
| D09 | -64.4 ± 0.4 | -79.8 ± 0.3 |
| D10 | -91.7 ± 0.4 | -81.4 ± 0.4 |

To identify the Bn-receptor complex that better meets the experimental information available, we analyzed the most important residues contributing to the binding free energy using the MMGBSA decomposition method [57], as explained in the methods section. Tables S1 and S2 of the supplementary material show the contributions to the binding free energy of diverse residues of the Bn-BB1 and Bn-BB2 complexes, respectively. Comparison of these Tables points to different patterns of interactions, suggesting that the peptide is bound in diverse poses. Accordingly, we focused on the contributions to the binding free energy of those residues known to be important for binding and/or activity from mutagenesis studies [7] and are listed in Tables 2 and 3. In the case of the BB2 receptor, Gln120(3.32), Arg287(6.58) and Arg308(7.39) were demonstrated to be key for Bn binding [17–19], along with residues Pro198(45.52), Ala307(7.38) and Phe184(ECL2) that were identified as binding modulators [18–21].

**Table 2.** Residue contributions to the binding free energy computed by means of the MMPBSA method for the bombesin-BB1 from the last 20 ns of the 200 ns relaxation MD trajectories (D01-D10).

| REPLICATES | $\Delta G$ CONTRIBUTION PER-RESIDUE (kcal/mol) | | | | |
| --- | --- | --- | --- | --- | --- |
|  | GLN<sup>3.32</sup> | ILE<sup>ECL2</sup> | PRO<sup>45.52</sup> | ARG<sup>6.58</sup> | ARG<sup>7.39</sup> |
| D01 | -0.65 ± 0.02 | -3.29 ± 0.01 | -0.33 ± 4.1E-3 | -4.13 ± 0.03 | -0.06 ± 0.01 |
| D02 | -1.35 ± 0.02 | -1.19 ± 0.01 | -0.32 ± 1.7E-3 | -1.92 ± 0.02 | -1.28 ± 0.01 |
| D03 | 0.12 ± 7.1E-4 | -3.31 ± 0.01 | -1.04 ± 4.5E-3 | -2.39 ± 0.03 | -1.82 ± 0.02 |
| D04 | -2.26 ± 0.02 | -1.98 ± 0.01 | -1.47 ± 0.01 | -5.69 ± 0.02 | -2.95 ± 0.02 |
| D05 | -0.43 ± 0.01 | -3.21 ± 0.02 | -1.18 ± 0.01 | -0.24 ± 0.02 | -0.65 ± 4.6E-3 |
| D06 | -0.30 ± 0.01 | -2.00 ± 0.01 | -2.64 ± 0.01 | -1.41 ± 0.02 | 0.07 ± 3.9E-4 |
| D07 | -0.06 ± 5.0E-3 | -1.99 ± 0.01 | -0.91 ± 0.01 | -0.93 ± 0.02 | -1.77 ± 0.01 |
| D08 | -3.69 ± 0.03 | -2.43 ± 0.01 | -1.58 ± 0.01 | -1.22 ± 0.01 | -0.37 ± 0.01 |
| D09 | -0.13 ± 0.01 | -2.70 ± 0.01 | -3.65 ± 0.01 | -0.87 ± 0.02 | -2.84 ± 0.02 |
| D10 | 0.08 ± 4.3E-4 | -1.75 ± 0.01 | -1.81 ± 0.01 | -3.79 ± 0.02 | -1.49 ± 0.01 |

Inspection of Table 3 reveals that complexes D01, D03, D06 and D07 show these residues with important contributions to the binding free energy. Interestingly, these four complexes are among those with the highest ΔG_bind_, as shown in Table 1. Inspection of the corresponding structures reveals that only D01, D06 and D07 exhibit Met14 - known to be important for Bn activity-sitting in the proximity of Trp277(6.48), which forms part of the highly conserved CWxP motif, involved in the activation class A GPCRs [59]. Finally, to discriminate among the three structures, we hypothesized that since the side chains of Trp8 and His12 do not contribute much to the binding free energy, it could be considered that they may rather play a role in the stabilization of the secondary structure of the peptide. Inspection of the three structures shows that only structure D07 exhibits the side chains of both residues at a close distance to each other, playing a putative role of providing robustness to the helical section of the peptide bound to the receptor. After, these considerations conformation D07 was selected as the most likely model for the Bn-BB2 complex, which is the second-best complex in energy terms (Table 1).

**Table 3.** Residue contributions to the binding free energy computed by means of the MMPBSA method for the bombesin-BB2 complex from the last 20 ns of the 200 ns relaxation MD trajectories (D01-D10).

| REPLICATES | $\Delta G$ CONTRIBUTION PER-RESIDUE (kcal/mol) | | | | |
| --- | --- | --- | --- | --- | --- |
|  | GLN<sup>3,32</sup> | PHE<sup>ECL2</sup> | PRO<sup>45,52</sup> | ARG<sup>6,58</sup> | ARG<sup>7,39</sup> |
| D01 | -1.07 ± 0.01 | -2.06 ± 0.01 | -2.75 ± 0.01 | -0.39 ± 0.02 | -3.28 ± 0.02 |
| D02 | -2.90 ± 0.02 | -2.00 ± 0.02 | -0.91 ± 0.01 | 0.20 ± 0.01 | -1.97 ± 0.01 |
| D03 | -2.40 ± 0.02 | -1.76 ± 0.01 | -1.10 ± 0.01 | -0.51 ± 0.01 | -3.03 ± 0.02 |
| D04 | -1.94 ± 0.01 | -1.94 ± 0.01 | -2.24 ± 0.01 | 0.42 ± 0.02 | -4.95 ± 0.02 |
| D05 | -0.04 ± 0.01 | -2.80 ± 0.01 | -1.40 ± 0.01 | -1.53 ± 0.02 | -1.96 ± 0.03 |
| D06 | -0.44 ± 0.01 | -1.21 ± 0.01 | -2.34 ± 0.01 | -2.55 ± 0.02 | -0.77 ± 0.02 |
| D07 | -1.32 ± 0.01 | -1.87 ± 0.02 | -1.21 ± 0.01 | -6.10 ± 0.03 | -2.71 ± 0.02 |
| D08 | 0.02 ± 2.3E-3 | -1.68 ± 0.01 | -0.95 ± 0.01 | -2.63 ± 0.02 | -2.03 ± 0.01 |
| D09 | -0.37 ± 0.02 | -2.03 ± 0.01 | -3.14 ± 0.01 | -0.75 ± 0.01 | 0.27 ± 0.02 |
| D10 | -1.04 ± 0.01 | -2.49 ± 0.01 | -1.22 ± 0.01 | 0.03 ± 0.01 | -2.82 ± 0.02 |

In the case of the BB1 receptor, a smaller number of mutagenesis studies are available. However, since BB1 exhibits a high sequence identity with the BB2 receptor in the orthosteric site, unless the extracellular loops play a definitive role in dictating the conformation of the bound Bn, it is expected that the peptide binds in similar fashion to both receptors. Consequently, most of the residues shown to be key for Bn binding to the BB2 receptor should also be key for BB1, assumption that cannot be ruled out from the experimental results available. Accordingly, we also analyzed in this case the contributions of residues Gln123(3.32), Ile187(ECL2), Pro200(45.52), Arg289(6.58) and Arg310(7.39) to the binding free energy. The results listed in Table 2, indicate that only complexes D04 and D08 exhibit simultaneously these residues involved in complex formation. However, none of the two complexes exhibits the intramolecular interaction between Trp8 and His12 side chains. This made us consider that we may have not found a suitable complex Bn-BB1 within the ten SMD trials. Accordingly, we took the Bn bound to BB2 structure D07 and constructed a Bn-BB1 complex from it and check its viability. Next, the complexes Bn-BB1 (constructed from D07) and Bn-BB2 (D07) we subject of four 500ns MD replicas to investigate the robustness of the structures (Figure S7 and S8 of the supplementary material) [49].

The best complex of each receptor was subject of additional 2μs MD simulation to ensure a robust stabilization of the complexes. Time evolution of the binding free energy (ΔG_bind_) during the 2μs MD simulation for the complexes of Bn-BB1 and Bn-BB2, computed using the MMGBSA method [52] is shown in Figure 1. Inspection of the Figure reveals a robust convergence of the ΔG_bind_ in both systems, displaying the corresponding value for the Bn-BB2 complex a more negative value than that of Bn-BB1 complex all the time, in agreement with the differential affinity found experimentally. Despite minor energy fluctuations, the value of the interaction was computed as the average value from the last 100ns in each complex, being -101.6 ± 0.5 kcal/mol for Bn-BB1 and -106.3 ± 0.4 kcal/mol for Bn-BB2, providing a binding free energy of 4.7±0.5 kcal/mol that agrees well with the differential affinity found experimentally [12].

**Figure 1.**
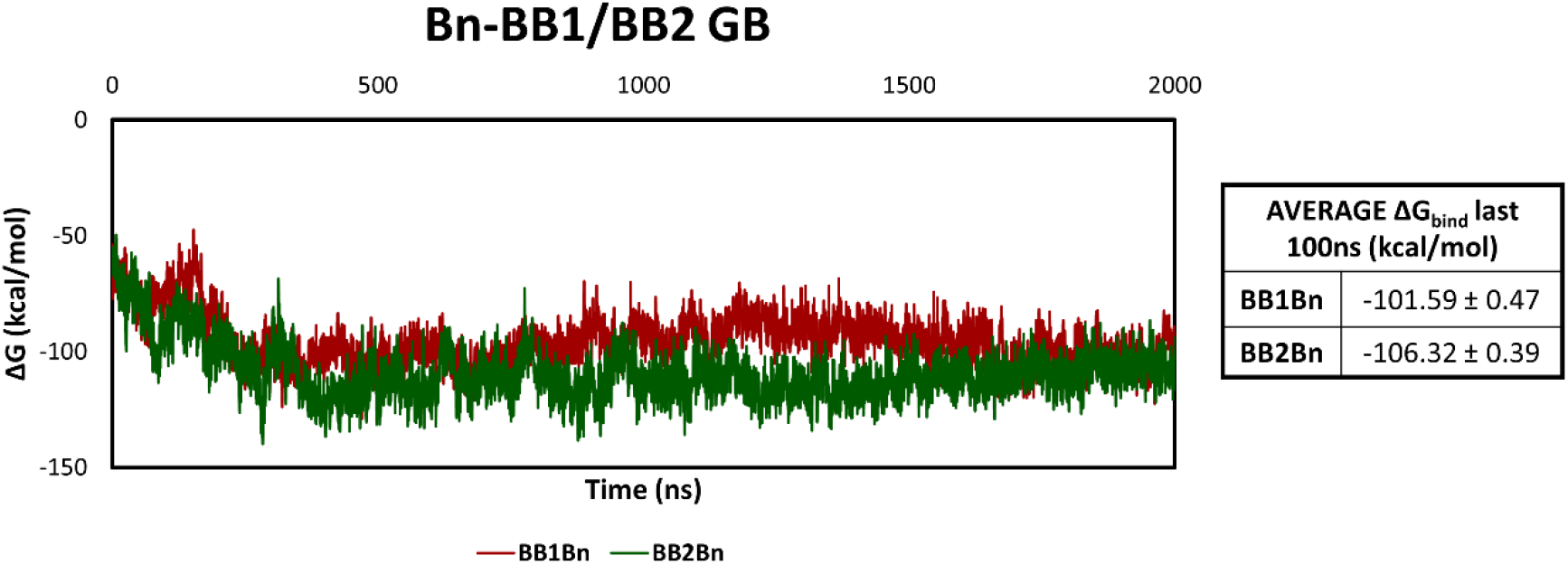
Time evolution of the binding free energy (ΔGbind) for the complexes of Bn-BB1 and Bn-BB2, computed by means of the MMGBSA method during the 2μs MD simulation.

### Structures of Bombesin Bound to the BB1 and BB2 receptors

In order to monitor the diverse conformations adopted by the complexes during the MD process, the two trajectories were subject of cluster analysis, as described in the methods section. For this purpose, we used 200,000 frames for each system, which corresponds to extracting one structure every 10ps. Moreover, sieve option equals to 4 was used in the clustering process, meaning that it uses 50000 frames and adds the other frames to the closest cluster in each case. Those residues with lower flexibility (RMSF < 0.5Å) were used to superimpose the structures sampled during the MD simulation. Subsequently, identification of the different structural features from ECL2 and ECL3 in Bn-BB1 and Bn-BB2 was performed by grouping similar structures in each complex into 10 different clusters using the average linkage algorithm [60], implemented in the *cpptraj* module of Amber18 [42, 51]. For this purpose, two clusterization processes were performed using alternatively the RMSD of the Cα from residues in ECL2 or the RMSD of the Cα from residues in ECL3 as distance for the clusterization process, for both Bn-BB1 and Bn-BB2 complexes.

The distribution of structures, as well as distances between members of the same cluster of the five most populated clusters and the two alternative clustering processes for both trajectories is shown in Tables S3 and S4 of the supplementary material, respectively. Inspection of the diverse clusters evidences large structural differences in the loops (Figure 2), being most significant in the ECL2. This loop exhibits a well-structured β-sheet motif conserved in other peptide activated GPCRs, however in contrast to the Bn-BB1 structure, in the Bn-BB2 it adopts a solvent exposed conformation. This, can be in part attributed to the tilting of the TM2 at its C-terminus in the BB2 model, compared with that in the BB1, which could also be considered as an artifact of the model. This differential feature also favors that some residues in TM2 of BB2 contribute to the binding free energy, as discussed below.

**Figure 2.**
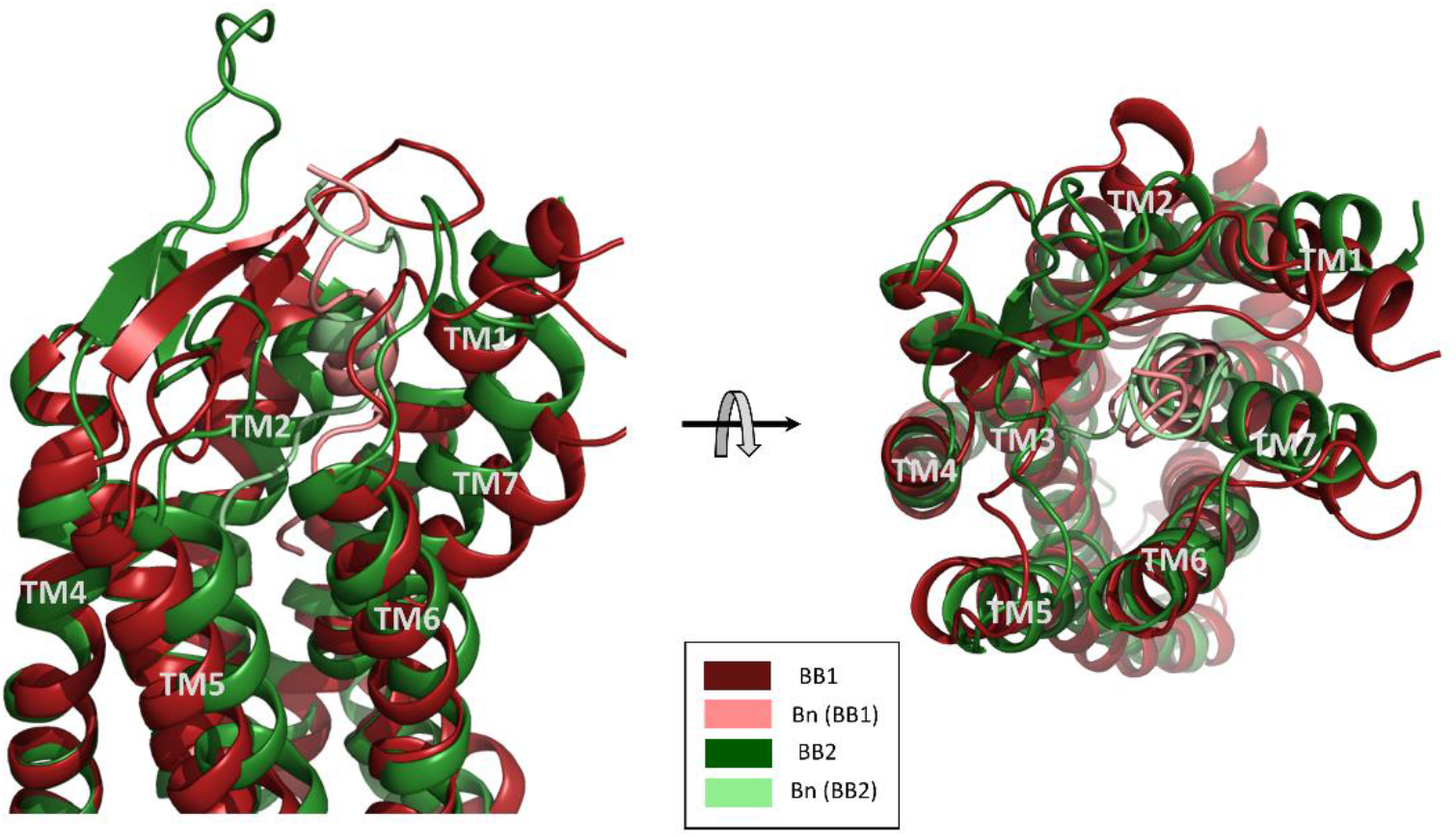
Superimposition of the most populated cluster for the two Bn-receptor complexes found in the two different cluster analysis.

Analysis of the RMSF of both complexes from the 2μs MD trajectory (Figures S9a and S9b of the supplementary material), shows that the Bn-BB2 complex exhibits higher number of residues with low fluctuations (<1Å) compared to the Bn-BB1 complex, suggesting a more robust structure. Interestingly, analysis of the fluctuations per residue reveals important differences between the two receptors, focusing on the ECLs. Figure 3 shows the comparison of the RMSF per residue between Bn-BB1 and Bn-BB2 during the 2μs MD, showing that ECL2 in BB2 exhibits higher fluctuations, in agreement with the perception that the loop is more solvent exposed than in the Bn-BB1 complex. Moreover, ECL3 also exhibits a differential behavior that can be attributed to the stereochemical features of the non-conserved residue Pro(ECL3) in BB1 versus Thr(ECL3) in BB2. This difference produces not only the change in the direction of the peptide backbone in BB1 due to the restricted conformational features of proline, but in BB2 threonine stablishes an interaction with bombesin Gln7, yielding as result an increased binding free energy, as discussed below.

**Figure 3.**
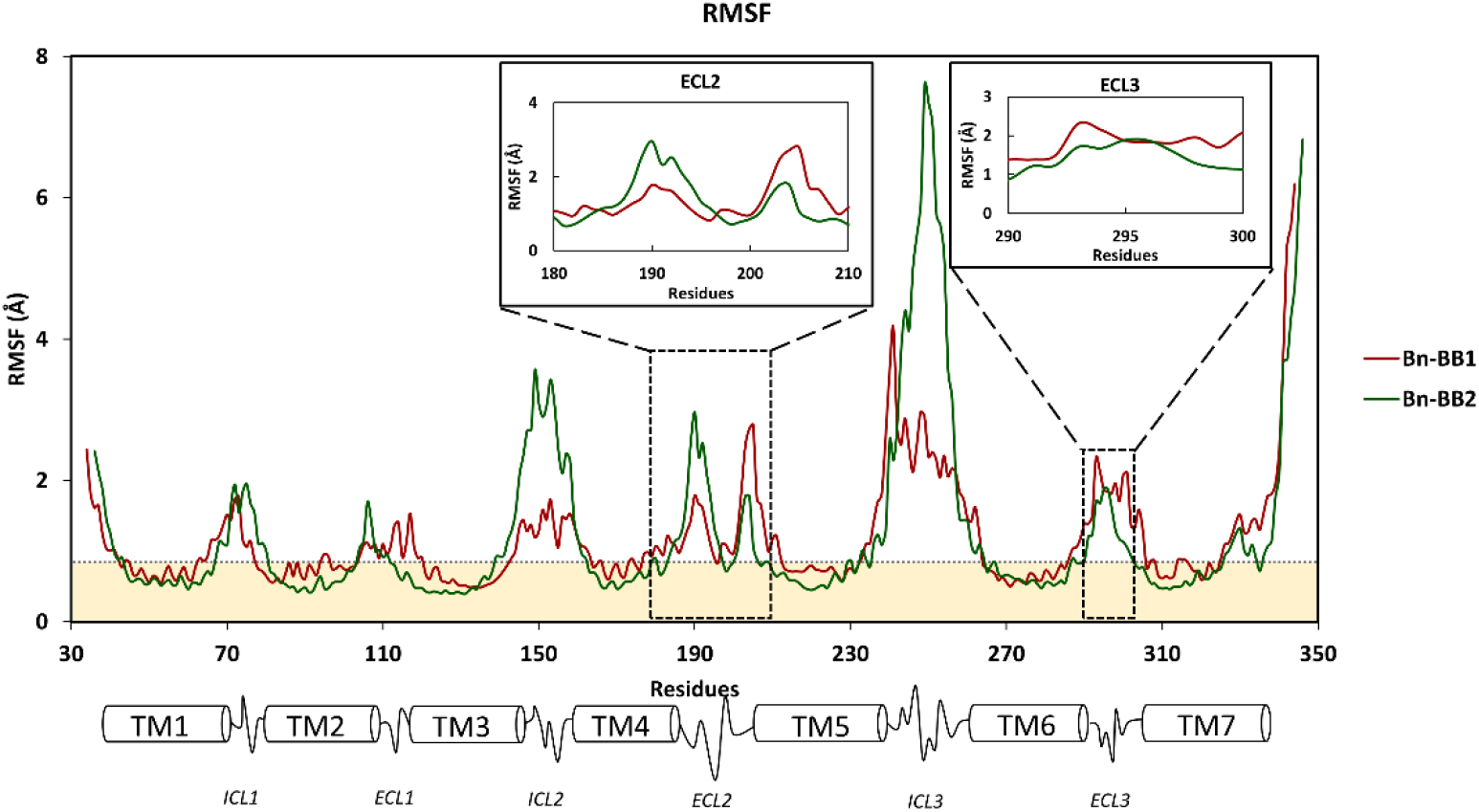
Comparison of the root-mean-square fluctuations (RMSF) per residue of Bn-BB1 and Bn-BB2 complexes during the 2μs MD simulation.

The differential binding free energy observed between the two complexes can be traced back to the contributions made by the most relevant residues involved in binding, computed during the last 100ns of each system and listed in Table 4. On the one hand, the energy contributions of Gln3.32, Ile/Phe(ECL2) and Pro45.52 are important as determined by site directed mutagenesis. Moreover, Arg7.39 also exhibits a relevant contribution to the binding free energy that it is expected to be similar in both receptors. However, inspection of Table 4 reveals that the contribution in Bn-BB1 is more relevant than in Bn-BB2.

**Table 4.** Contribution to the binding free energy of a few relevant residues computed by means of the MMPBSA method during the last 100 ns of the 2 μs trajectory for the Bn-BB1 and Bn-BB2 complex.

| Residue | Bn-BB1 | Bn-BB2 |
| --- | --- | --- |
| GLN<sup>3.32</sup> | $-2.21 \pm 4.5\text{e-}3$ | $-2.05 \pm 0.01$ |
| ILE/PHE<sup>ECL2</sup> | $-2.43 \pm 3.9\text{e-}3$ | $-3.79 \pm 0.01$ |
| PRO<sup>45.52</sup> | $-2.44 \pm 3.9\text{e-}3$ | $-3.19 \pm 4.3\text{e-}3$ |
| ARG<sup>6.58</sup> | $-0.62 \pm 4.5\text{e-}3$ | $-0.97 \pm 0.01$ |
| ARG<sup>7.39</sup> | $-6.19 \pm 0.01$ | $-3.04 \pm 0.01$ |

To find an explanation, we inspected of the relative orientation of the residue relative to the peptide, resulting in that the arginine interacts with the backbone of Leu13 in a similar manner in the two complexes (see Figure S10 of the supplementary material). Then, pairwise contributions to the binding free energy from the peptide perspective side were analyzed (Table 5), showing that the interaction energy between the two residues is the same for both receptors (-7.04 ± 0.01 kcal/mol in Bn-BB1 and -6.65 ± 0.01 kcal/mol in Bn-BB2). The reason for the differential energy contribution observed in Table 4, is actually produced by the proximity of Val10 to Arg7.39 in BB1, but not in BB2 (Figure S10 of the supplementary material) that supplies an additional interaction through its backbone and contributes to a higher binding free energy for Arg7.39 in the decomposition process. Accordingly, the interaction of Arg7.39 with the peptide backbone can be considered as similar in both receptors. In regard to residue Arg6.58, despite experimental findings suggesting an important role in Bn binding, present results show only a poor contribution to the binding free energy (<-1 kcal/mol) (see Figures 5A and 5B). In order to understand this discrepancy, we investigated if the solvent could mediate the interaction. Accordingly, we looked for water molecules in the neighborhood of Arg6.58 and computed again the binding energy of the residue.

**Table 5.** Bombesin (Bn) residue contributions to the binding free energy computed by mean of the MMPBSA method using the last 100 ns of the 2 μs trajectory. The intramolecular interaction His12…Trp8 in bombesin is also listed at the end of the Table.

| Bn Residues | BB1 Residues | $\Delta G$ Contribution (kcal/mol) | BB2 Residues | $\Delta G$ Contribution (kcal/mol) |
| --- | --- | --- | --- | --- |
| GLN 7 | ILE<sup>ECL3</sup> | $-3.01 \pm 0.02$ | THR<sup>ECL3</sup> | $-4.01 \pm 0.02$ |
| TRP 8 | PRO<sup>45.52</sup> | $-1.80 \pm 0.01$ | PRO<sup>45.52</sup> | $-3.48 \pm 0.01$ |
| | ARG<sup>6.58</sup> | $-2.99 \pm 0.02$ | ARG<sup>6.58</sup> | $-2.00 \pm 0.02$ |
| VAL 10 | ARG<sup>6.58</sup> | -0.58 ± 3.9E-3 | ARG<sup>6.58</sup> | -1.09 ± 0.01 |
|  | ARG<sup>7.39</sup> | -5.85 ± 0.01 | ARG<sup>7.39</sup> | -0.83 ± 1.5E-3 |
| LEU 13 | PRO<sup>3.29</sup> | -1.18 ± 4.2E-3 | PRO<sup>3.29</sup> | -1.55 ± 0.01 |
|  | GLN<sup>3.32</sup> | -2.37 ± 0.01 | GLN<sup>3.32</sup> | -1.75 ± 0.02 |
|  | ARG<sup>7.39</sup> | -7.04 ± 0.01 | ARG<sup>7.39</sup> | -6.65 ± 0.01 |
| MET 14 | GLN<sup>3.32</sup> | -2.39 ± 0.01 | GLN<sup>3.32</sup> | -2.92 ± 0.01 |
| **HIS 12** | **TRP 8** | -1.41 ± 0.01 |  | -1.76 ± 0.01 |

Interestingly, the result is significantly higher (-5.86 ± 0.02 kcal/mol) in Bn-BB2, compared to the result on the Bn-BB1 complex (-2.42 ± 0.02 kcal/mol) (Figure 4A and 4B). This suggests that the interaction is relevant for binding as found experimentally and larger for the Bn-BB2 complex. Finally, two more residues not listed in Table 4 also exhibit a higher contribution to the binding energy in BB2 compared to BB1. This includes residues Arg2.64 in TM2 and Thr(ECL3). In the case of Arg2.64, the difference is due to the differential conformation adopted by TM2 in both receptors; while in BB1 is in a more open and distant from Bn, in BB2 is adjacent to Bn and engaged in an interaction with Ala9 (Figure S11 of the supplementary material). On the other hand, Thr(ECL3) is closer to Bn in BB2, interacting with Gln7. This interaction is prevented by the presence of Pro(ECL3) in BB1 that renders a flexibility difference in ECL3 and in addition, lacks a polar group in its sidechain.

**Figure 4.**
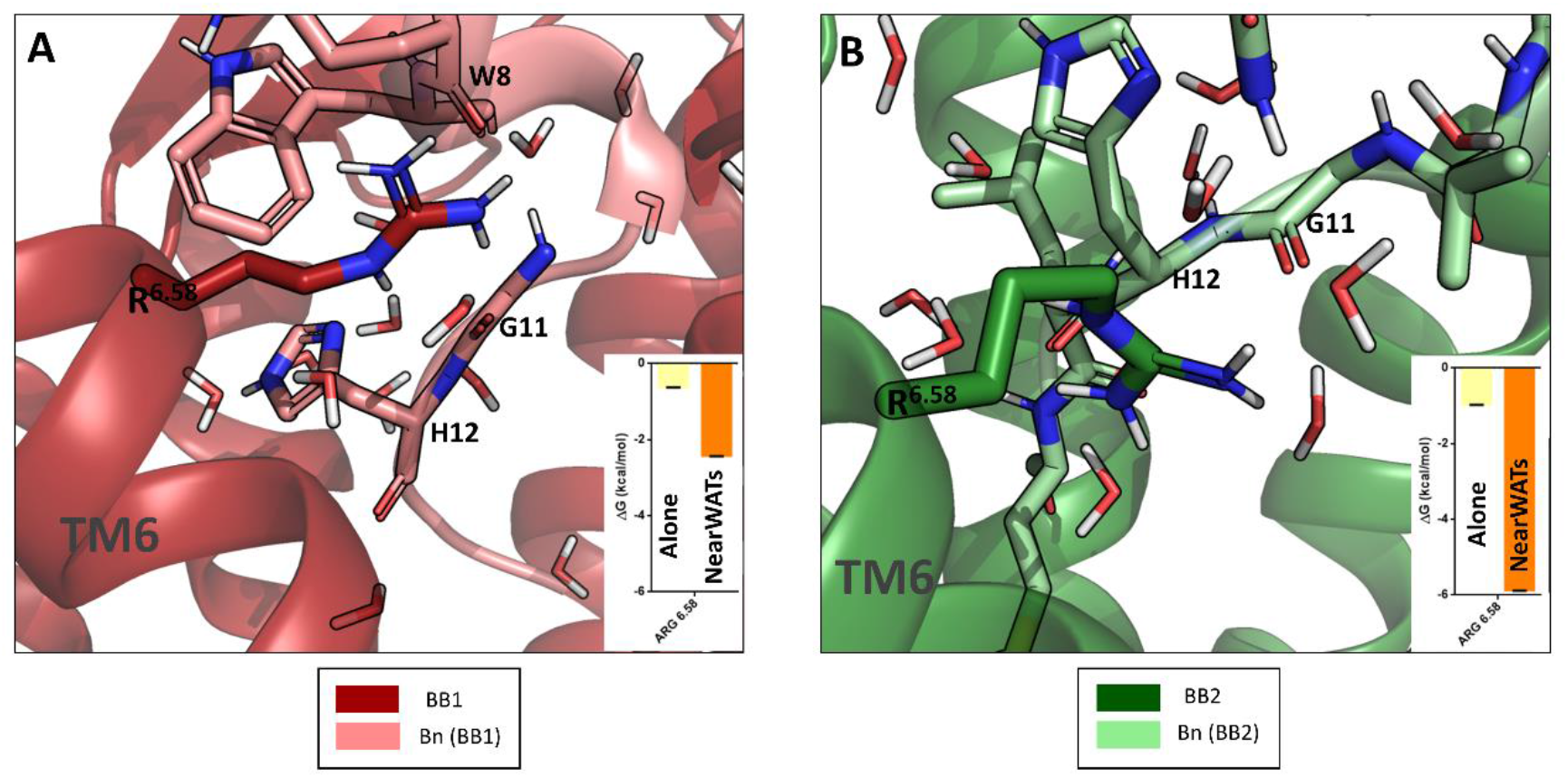
Differential interaction of residue Arg6.58 and the peptide in the two receptors. A) Bn-BB1 complex; B) Bn-BB2 complex.

The analysis of the peptide-receptor interactions from the peptide perspective, are listed for both receptors in Table 5. The most relevant residues of bombesin C-terminal for binding are Trp8, His12, Leu13 and Met14. As has been mentioned previously, Met14 is also key for receptor activation, since its deletion transforms the peptide into a potent antagonist [14–16]. As can be seen, contributions from these residues are similar in both complexes. Specifically, Trp8, Leu13 and Met14 are those that contribute the most to the binding. In contrast, His12 is not as relevant in energy terms as the other three (-0.53 ± 0.01 kcal/mol and -2.07 ± 0.01 kcal/mol for the Bn-BB1 and Bn-BB2 complexes, respectively). This provides further support to consider that His12 may not have a direct role in binding, but its role may be structural, to keep the helical conformation from bombesin through an intermolecular interaction with Trp8 that occurs in both systems. Despite the similar patterns in the free energy decomposition, it is important to underline that most of the peptide residues interact with the same residues in BB1 and BB2. Thus, Trp8 interacts mainly with Pro45.52 and Arg6.58; Leu13 with Pro3.29, Gln3.32 and Arg7.39 and Met14 with Gln3.32 in both receptors (Table 5). In contrast, His12 interacts with Arg6.58 in BB2, but not in BB1, and Gln7 interacts with a Thr(ECL3) in BB2 but not in BB1, since there the residue is a Pro. This difference is also connected to the flexibility of ECL3 in BB2, contributing to the differential affinity of bombesin to the two receptors.

Hydrogen bonds taking place during the last 100ns of MD trajectory for each complex are listed in Table S5 of the supplementary material. Analysis of the Table points to a conserved interaction between the Arg7.39 and Leu13 (Figure S10 of the supplementary material). In both complexes, the lifespan of this hydrogen bond is above 80% (89.1% for Bn-BB1 and 86.6% for Bn-BB2). Moreover, the other primary amine of the arginine is also engaged in hydrogen bonding, although the lifespan for this second hydrogen bond donor is shorter, but still significant occurrence (56.6% for Bn-BB1 and 40.3% for Bn-BB2). Furthermore, there is a common interaction between Ala9 and TM2 in both receptors (Figure S11 of the supplementary material). In the Bn-BB1 complex, this hydrogen bond occurs between Ala9 and Tyr2.65 (54.1%); and in Bn-BB2, the same hydrogen bond is found (56.8%), but also a small percentage hydrogen bond between Ala9 and Arg2.64 is observed (19.7%). However, despite this similarity, there are some noteworthy differences. The most relevant difference involves Arg6.58. This residue forms hydrogen bonds with water molecules near the Bn-peptide in BB2 but in a lesser extend in the BB1 complex (Figure S12 of the supplementary material). This differential interaction renders an explanation for the energy contribution of Arg6.58 to the binding free energy.

In order to get a better understanding of the role of loops in the structure of the bound conformation, a few distances between residues of the loops and Bn were monitored during the 2μs MD. Figure 5 shows the time evolution of the distance between ECL3 and ECL2 and between ECL2 or ECL3 with Bn. In the Bn-BB1 system, distances change dramatically after the first 200ns, remaining stable for the rest of the trajectory with the N-terminal of ECL3 close to both, ECL2 and Bn. On the other hand, the distance between the C-terminal of ECL3 and ECL2 remains constant, showing small fluctuations, whereas the distance with Bn peptide increases and between ECL2 and Bn peptide decreases. This suggests that in BB1, Bn accommodates adjacent to the ECL2 and moves away from ECL3. In contrast, in the Bn-BB2 system more coordinated movements are observed since in appears that changes occur in a coordinated fashion. Comparison between the two systems shows that the distance between the C-terminal of the ECL3 and Bn is shorter in the Bn-BB2 system, favoring receptor-peptide interactions, confirming the results of the per-residue binding free energy decomposition, where residues of the loop interact with Bn in BB2, but not in BB1. In other words, Pro(ECL3) in BB1 produces a change in the direction of the amino acid chain that forces diverse residues of ECL3 to remain closer to the helix bundle. This in turn makes that relevant amino acids for Bn interaction such as Arg6.58, are prevented to adopt a proper conformation to interact with Bn. This can be seen in Figure 6, where the distance between Ile(ECL3) in BB1 and its homologous Val(ECL3) in BB2 with the Arg6.58 is shown. In this case, Arg6.58 in BB1 adopts a different conformation compared to BB2; whereas in BB2 this residue is oriented towards the interior of the binding site and in close proximity to Bn, in BB1 it is oriented towards TM7 with a notable interaction energy difference. This structural difference may explain the relevance of ECL3 demonstrated experimentally to differentiate between BB1 and BB2 in the case of selective ligands [20].

**Figure 5.**
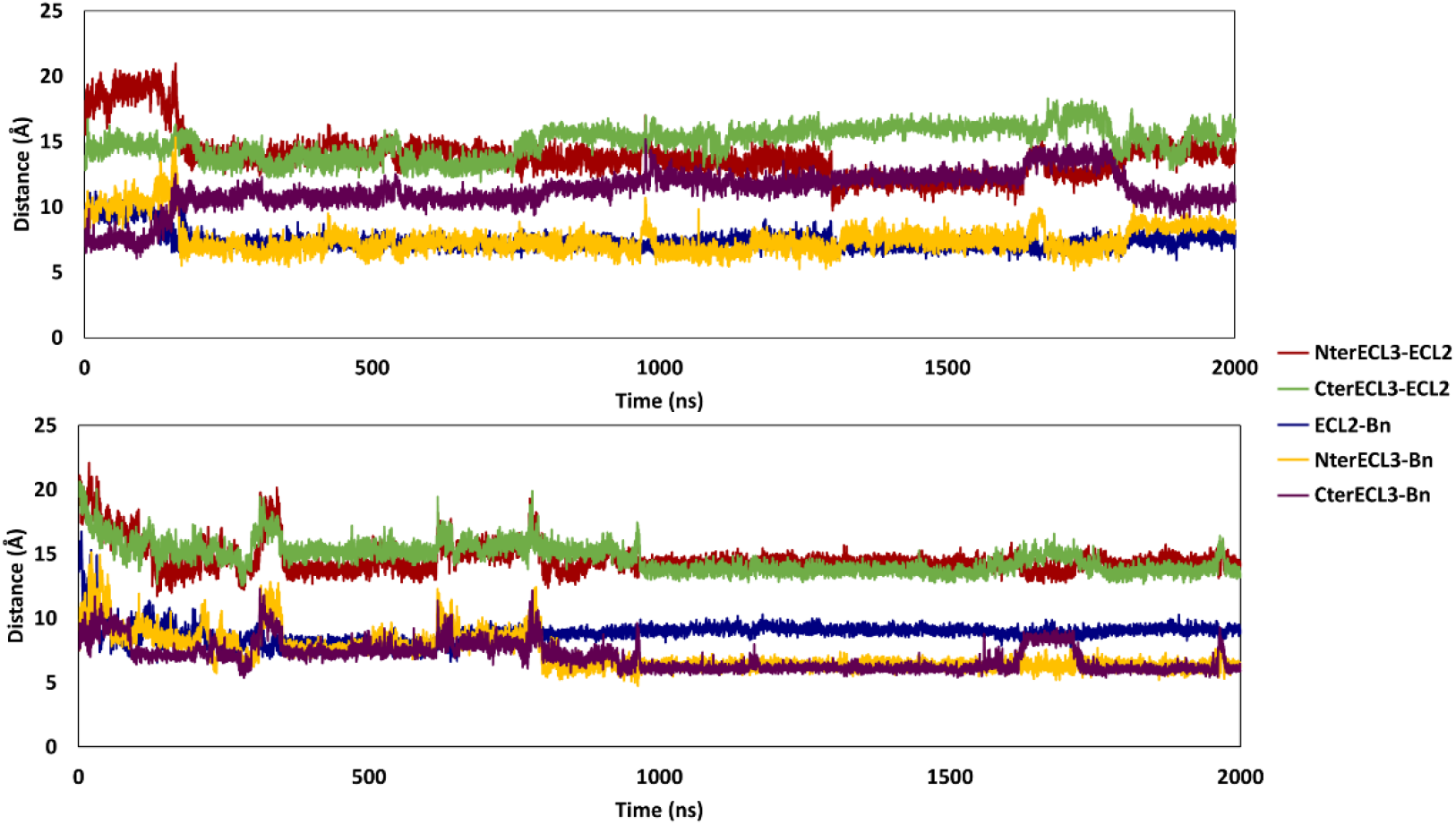
Time evolution of selected distances between ECL3 and ECL2, as well as that between ECL2 or ECL3 with Bn.

**Figure 6.**
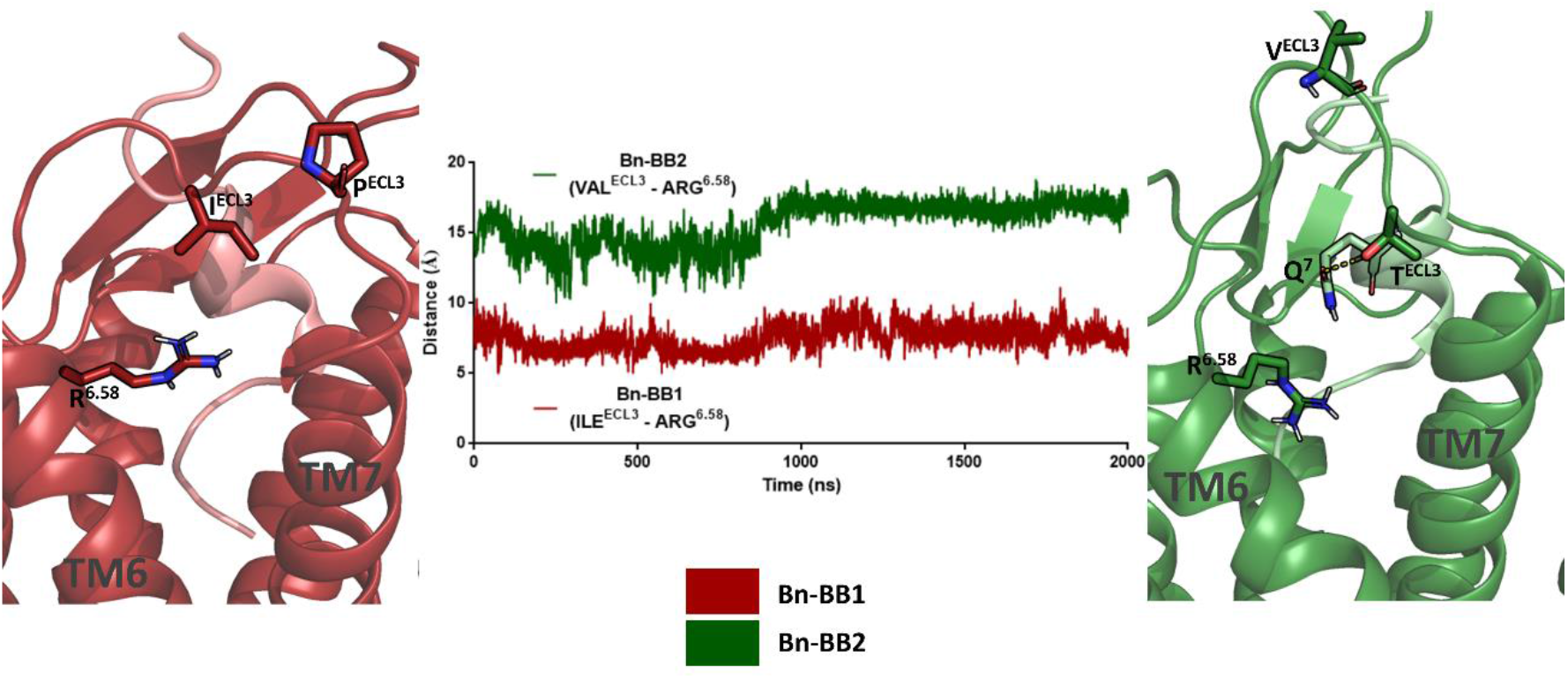
Distance between Ile(ECL3) in BB1 and its homologous Val(ECL3) in BB2 with the Arg6.58

## DISCUSSION

The present study describes the construction of 3D models of bombesin bound to the BB1 and BB2 receptors, respectively. For this purpose, rough models of the receptors were constructed by homology modeling using the endothelin B receptor as template and refined embedded in a lipid membrane, using molecular dynamics. The models were subsequently used for docking bombesin, by means of Steered Molecular Dynamics. For this purpose, the peptide in a helical structure, resulted from a previous computational study [31] was used as the starting geometry. Several trajectories were computed for any of the two receptors. After a careful analysis, the complex fulfilling best the available experimental results for each receptor was selected for further studies.

After 2μs MD trajectory the binding free energy of the systems was computed using the last 100ns by means of the MMPBSA method. The results show that the binding free energy is systematically lower for the Bn-BB2 complex along the simulation process, with an average difference with Bn-BB1 of 4.7±0.5 kcal/mol in the last 100ns that agrees well with the differential experimental affinity reported for the two receptors (K_i_=2 nM for BB1 and 0.07 nM for BB2). Analysis of the contributions to the binding energy per residue provides support to explain the effect of mutations on specific residues on the receptors or substitutions in the Bn sequence. A deeper analysis of these interactions permits to understand particularities that justify the differential affinity of Bn to the two receptors. Specifically, although there are a number of interactions that are similar in both complexes, a few of them are more favorable for the Bn-BB2 complex including Arg6.58 and Thr(ECL3) (a Pro in BB1). Moreover, this latter not conserved residue in ECL3 is responsible, in addition to provide an extra hydrogen bond interaction with the receptor, to prompt ECL3 to exhibit a differential dynamic behavior in the two receptors. Consequently, residue Arg6.58 in BB1 cannot adopt a suitable conformation for interacting with Bn, key interaction for the bound conformation of the peptide in BB2.

This study on the bound conformation of bombesin to BB1 and BB2, allowed us to identify the main features involved in binding, elucidating the differences that explain the different affinity observed for both receptors. Furthermore, these results shed light into the interaction requirements needed for the activation of bombesin receptors, being a reference point for designing future experiments.

## METHODS AND MODELS

### Construction and Refinement of the BB1 and BB2 Receptors by Homology Modeling

Atomistic models of the BB1 and BB2 receptors were constructed by homology modelling using MODELLER v9.22.22 [35]. For this purpose the human endothelin B receptor (PDB entry code 5X93) [36], with a 38% of sequence similarity to BB1 and 36% to BB2 receptors was used as the template. From the diverse models constructed for each receptor, those with the highest DOPE score [37] were used for subsequent refinement through molecular dynamics (MD) [38]. In order to avoid artifacts during the construction process, the N-terminus was removed in both models. To carry out the MD simulations, receptors were embedded in a lipid bilayer composed of 1-palmitoyl-2-oleoyl-sn-glycero-3-phosphocholine (POPC) and water molecules. Systems were generated according to the procedure described previously [39, 40] by means of PACKMOL-Memgen [41]. Next, a set of randomly selected water molecules were replaced by sodium and chloride ions to have a neutral system with a concentration of 0.15 M of sodium chloride. Subsequently, the LEaP module from Amber18 [42] was used to prepare each system to perform MD calculations using the ff14SB, LIPID17 and TIP3P force fields [43, 44]. Equilibration of the systems was performed as described elsewhere [45, 46] and electrostatic interactions were treated using the Particle Mesh Ewald procedure [47]. Three 500ns replicas were performed for each system at 300K using the Langevin thermostat [48] to keep the temperature constant. Multiple production runs were used to enhance conformational sampling and to ensure energy convergence [49]. MD calculations were performed under the isothermic-isobaric ensemble (NPT), fixing all bonds involving hydrogen atoms with the SHAKE algorithm [50] allowing us to use a time step of 2 fs. The most frequently sampled structures for apoBB1 and apoBB2 systems were used as final receptor models, respectively. For this purpose, the structures sampled during the MD refinement process were subjected to cluster analysis using the average linkage algorithm, implemented in the *cpptraj* module of Amber18 [42, 51]. The root mean square deviation (rmsd) of the Cα of residues belonging to the transmembrane domain (TM1-7] was used as a measure of the distance between two structures. Final models of the two receptors corresponded to the representative structures of the most populated clusters, respectively.

### Steered Molecular Dynamics

SMD calculations on each of the refined models of the BB1 or BB2 receptors, embedded in a lipid bilayer and Bn in its prospective conformation in solution taken from a previous computational study [31], were run in parallel. The starting configuration corresponds to the peptide and the protein at a distance long enough from each other to ensure the complete solvation of both molecules (Figure 7). SMD calculations were performed using a constant external force to pull the peptide with its C-terminus facing the receptor extracellular side, towards the fixed receptor through a straight line. The distance between the peptide and the protein was used as pulling variable of choice that was computed using reference points on both, the ligand and the receptor, respectively. Specifically, for the BB1 receptor the reference point was located at the geometrical center of residues Leu92(2.53), Val127(3.36), Trp279(6.48) and Asn316(7.45) and for the BB2 receptor, the geometrical center of residues Leu89(2.53), Val124(3.36), Trp277(6.48) and Phe312(7.43). On the other hand, the reference point on the peptide was located at the geometrical center of residues Gly11, His12 and Leu13. In the present work, the distance receptor-Bn was set approximately to 25Å for BB1 and 32Å for BB2, respectively. The difference is due to the different special distribution of the extracellular loops. SMD calculations ended when the distance reached a specified threshold value of 5Å, which corresponds to the C-terminus of the peptide accommodated in the orthosteric site of the receptor. Pulling of the peptide is performed in small steps, allowing the system to relax after each move. Different force constants and pulling velocities were investigated before production SMD trajectories (data not shown). Finally, ten independent 50ns SMD trajectories computed in steps of 0.002 ps were performed for each receptor, applying a force constant of 50 kcal/mol·Å^2^. After completion of each trajectory, structures were subsequently relaxed through a 200 ns MD calculation to accommodate the peptide bound to the receptor, using the last 20 ns to compute the binding free energy. Additional calculations were carried out to assess the effect of the starting side chains orientation on the final structures obtained after the relaxation process that followed the SMD calculations.

**Figure 7.**
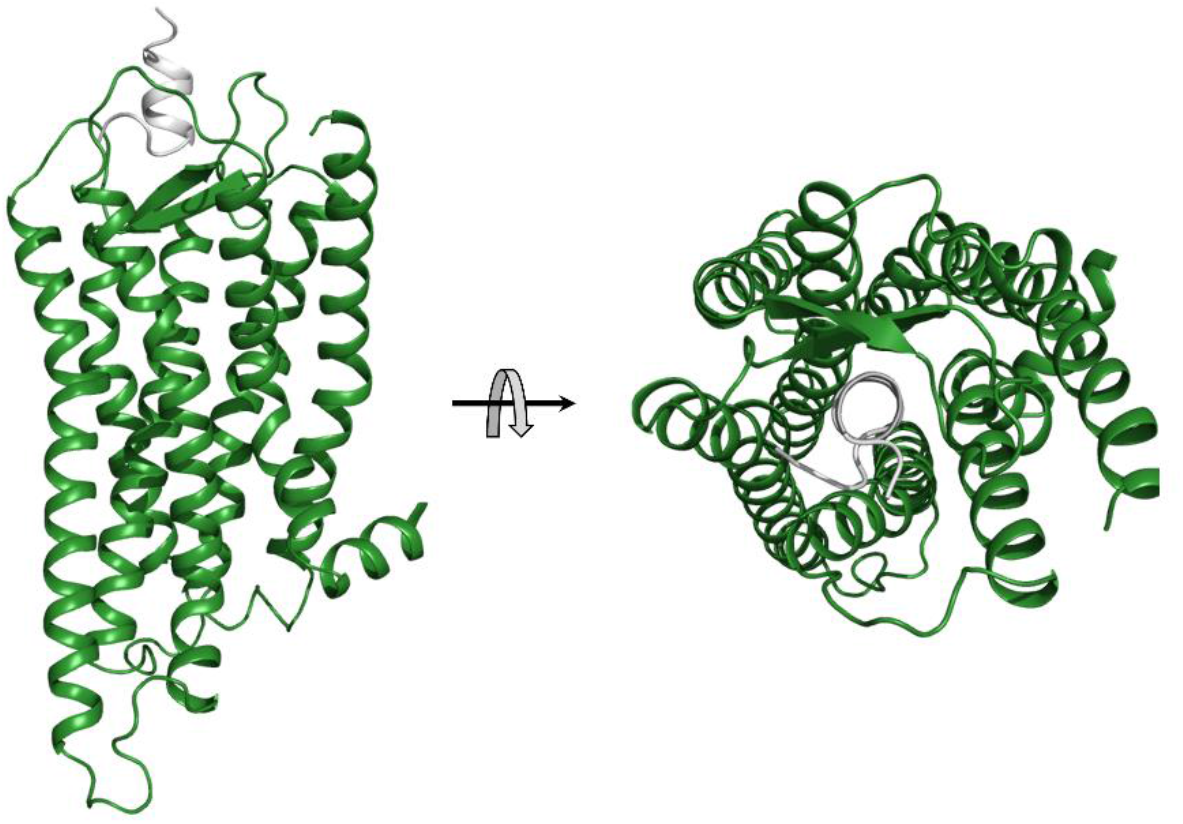
Starting configuration of the peptide (white) and the receptor (green) at a distance long enough from each other to ensure the complete solvation of both molecules.

### Binding Free Energy Calculations

Binding Free energies were computed using the Molecular Mechanics Generalized - Born Surface Area (MMGBSA) algorithm [52]. In this method, free binding energies are computed according to the equation:

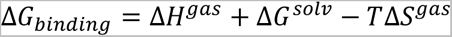

where ΔH^gas^ is the gas-phase interaction energy calculated by summing the internal energy, the noncovalent van der Waals, and electrostatic molecular mechanics energies. On the other hand, ΔG^solv^ is computed as the result of the sum between polar and non-polar terms. The former can be estimated numerically using the Generalized Born (GB) method [53]. In the present work, we used the Onufriev-Bashford-Case (OBC) generalized Born method (igb=2) [54]. Concerning the latter, it can be calculated using the following equation:

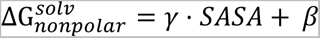

where SASA is the Solvent-Accessible Surface Area, calculated using the linear combinations of pairwise overlaps (LCPO) method [55] and the values for γ and β constants were set to 0.0072 kcal/mol·Å^2^ and 0.0 kcal/mol for MMGBSA [56]. Final values were averaged over 20 ns after ensuring that each of these trajectories was stable.

### Free Energy Decomposition

Binding free energy contributions per residue were computed using the snapshots of the last nanoseconds of each MD trajectory for each complex. They were computed using the MMGBSA decomposition method [57]. The procedure was carried out using the GB approach to calculate the electrostatic component of the solvation energy and the LCPO for the nonpolar component.

### Analysis of Hydrogen Bonds

Hydrogen bond occurrence between receptor residues and the peptide were analyzed using the *cpptraj* module of Amber18 [42, 51] and monitorized over the last 100ns of the simulation time. The criteria used for hydrogen bonding were to exhibit a distance between the hydrogen-bond acceptor and donor lower than 3.0Å and with the acceptor-H-donor angle, higher than 135°. Moreover, not only hydrogen bonds between BB1/BB2 and Bn peptide were considered, but also residues from both receptors and from the peptide with the nearest water molecules, following the same criterion.

### Distances

Distance between diverse residues were computed using the *cpptraj* module from Amber18 [42, 51]. Distances between the Cα of the residues Ile187(ECL2), Asn294 (NterECL3), Pro298 (CterECL3) and Leu4 (Bn) for the Bn-BB1 complex, and Phe184 (ECL2], Ser292 (NterECL3], Thr296 (CterECL3) and Leu4(Bn) for the Bn-BB2 complex. On the other hand, distances between ECL3 and Arg(6.58) were computed using different atoms of the lateral chains of Arg(6.58) and Ile(ECL3) and Val(ECL3) from Bn-BB1 and Bn-BB2 complexes respectively.

## ACKNOWLEDGEMENTS

This study was supported by the Agència de Gestió d’Ajuts Universitaris i de Recerca (AGAUR)-Generalitat de Catalunya (2021SGR00350 and 2021SGR00342), and Spanish Structures and Excellence María de Maeztu program, grant number CEX2021-001202-M.

## DATA AVAILABILITY

All input data, results from the molecular dynamics simulations and analysis is available on a GitHub repository at https://github.com/JaimeRubioMartinez/BnR_Bn.

## SUPPLEMENTARY INFORMATION

**Table S1.** Contributions to the binding free energy of diverse residues in the Bn-BB1 complex, computed by means of the MMPBSA method using the last 20 ns of the 200ns relaxation MD trajectory.

**Table S2.** Contributions to the binding free energy of diverse residues in the Bn-BB2 complex, computed by means of the MMPBSA method using the last 20 ns of the 200ns relaxation MD trajectory.

**Table S3.** Distances between members of the same cluster of the five most populated clusters using RMSD of the Cα from residues in ECL2.

**Table S4.** Distances between members of the same cluster of the five most populated clusters using RMSD of the Cα from residues in ECL3.

**Table S5.** Hydrogen bonds taking place during the last 100ns of the 2 μs MD trajectory for each complex.

**Figure S1.** Sequence alignment of the bombesin BB1 and BB2 receptors with the template endothelin B.

**Figure S2.** Time evolution of the root-mean-square deviation (rmsd) of the bombesin BB1 and BB2 receptors during the refinement process.

**Figure S3.** Time evolution of the root-mean-square deviation (rmsd) of the 200 ns trajectory of the bombesin-BB1 receptor complexes for the 10 (D1-D10) trials of Steered Molecular Dynamics.

**Figure S4.** Time evolution of the root-mean-square deviation (rmsd) of the 200 ns trajectory of the bombesin-BB2 receptor complexes for the 10 (D1-D10) trials of Steered Molecular Dynamics.

**Figure S5.** Time evolution of the binding free energy computed by means of the MMPBSA method for the 200 ns trajectory of the bombesin-BB1 receptor complexes for the 10 (D1-D10) trials of Steered Molecular Dynamics.

**Figure S6.** Time evolution of the binding free energy computed by means of the MMPBSA method for the 200 ns trajectory of the bombesin-BB2 receptor complexes for the 10 (D1-D10) trials of Steered Molecular Dynamics.

**Figure S7.** Time evolution of the binding free energy computed by means of the MMPBSA method for the complex bombesin-BB1.

**Figure S8.** Time evolution of the binding free energy computed by means of the MMPBSA method for the complex bombesin-BB2.

**Figure S9.** RMSF per residue for the bombesion-receptor complexes during the 2μs MD trajectory. A) RMSF for the Bn-BB1 complex; B) RMSF for the Bn-BB2 complex.

**Figure S10.** Relative orientation of the residue Arg7.39 interaction with Leu13 and differential position of Val10 in the Bn-BB1 and Bn-BB2 complexes.

**Figure S11.** Differential orientation of Arg2.64 and interaction with Ala9 in the two bombesin-receptor complexes.

## REFERENCES

1. Anastasi A, Erspamer V, Bucci M. Isolation and structure of bombesin and alytesin, 2 analogous active peptides from the skin of the European amphibians Bombina and Alytes. Experientia. 1971 Feb;27(2):166–7.

2. Erspamer V, Melchiorri P. Active polypeptides: from amphibian skin to gastrointestinal tract and brain of mammals. Trends Pharmacol Sci. 1980;1(2):391–5.

3. Jensen RT, Moody TW. Bombesin peptides (cancer). In: A.J. Kastin, editor. Handbook of Biologically Active Peptides. 2nd ed. Amsterdam; 2013. p. 506–11.

4. Märki W, Brown M, Rivier JE. Bombesin analogs: Effects on thermoregulation and glucose metabolism. Peptides. 1981 Jan 1;2(SUPPL. 2):169–77.

5. Mahmoud S, Staley J, Taylor J, Bogden A, Moreau JP, Coy D, et al. [Psi13,14] Bombesin Analogues Inhibit Growth of Small Cell Lung Cancer in Vitro and in Vivo1 | Cancer Research | American Association for Cancer Research. Cancer Res. 1991;51(7):1798–802.

6. Weber HC. Regulation and signaling of human bombesin receptors and their biological effects. Curr Opin Endocrinol Diabetes Obes. 2009 Feb;16(1):66–71.

7. Jensen RT, Battey JF, Spindel ER, Benya R V. International Union of Pharmacology. LXVIII. Mammalian bombesin receptors: nomenclature, distribution, pharmacology, signaling, and functions in normal and disease states. Pharmacol Rev. 2008 Mar;60(1):1–42.

8. Benya R V, Kusui T, Pradhan TK, Battey JF, Jensen RT. Expression and characterization of cloned human bombesin receptors. Mol Pharmacol. 1995;47(1).

9. Pooja D, Gunukula A, Gupta N, Adams DJ, Kulhari H. Bombesin receptors as potential targets for anticancer drug delivery and imaging. Int J Biochem Cell Biol. 2019;114(February):105567.

10. Perez J, Corcho F, Llorens O. Molecular Modeling in the Design of Peptidomimetics and Peptide Surrogates. Curr Med Chem. 2012 Oct 30;9(24):2209–29.

11. Perez JJ. Designing Peptidomimetics. Curr Top Med Chem. 2018 May 23;18(7):566–90.

12. Lin JT, Coy DH, Mantey SA, Jensen RT. Comparison of the peptide structural requirements for high affinity interaction with bombesin receptors. Eur J Pharmacol. 1995 Dec 27;294(1):55–69.

13. Gargosky SE, Wallace JC, Upton FM, Ballard FJ. C-terminal bombesin sequence requirements for binding and effects on protein synthesis in Swiss 3T3 cells. Biochem J. 1987;247(2):427.

14. Rivier JE, Brown MR. Bombesin, Bombesin Analogues, and Related Peptides: Effects on Thermoregulation. Biochemistry. 1978;17(9):1766–71.

15. Palmioli A, Ceresa C, Tripodi F, La Ferla B, Nicolini G, Airoldi C. On-cell saturation transfer difference NMR study of Bombesin binding to GRP receptor. Bioorg Chem. 2020 Jun 1;99(April):103861.

16. Broccardo M, Erspamer GF, Melchiorri P, Negri L, Castiglione R DE. Relative Potency of Bombesin-like Peptides. Br J Pharmacol. 1975 Oct 1;55(2):221–7.

17. Donohue PJ, Sainz E, Akeson M, Kroog GS, Mantey SA, Battey JF, et al. An aspartate residue at the extracellular boundary of TMII and an arginine residue in TMVII of the gastrin-releasing peptide receptor interact to facilitate heterotrimeric G protein coupling. Biochemistry. 1999 Jul 20;38(29):9366–72.

18. Lin Y, Jian X, Lin Z, Kroog GS, Mantey S, T. Jensen R, et al. Two amino acids in the sixth transmembrane segment of the mouse gastrin-releasing peptide receptor are important for receptor activation. J Pharmacol Exp Ther. 2000;294(3):1053–62.

19. Nakagawa T, Hocart SJ, Schumann M, Tapia JA, Mantey SA, Coy DH, et al. Identification of key amino acids in the gastrin-releasing peptide receptor (GRPR) responsible for high affinity binding of gastrin-releasing peptide (GRP). Biochem Pharmacol. 2005 Feb 15;69(4):579–93.

20. Tokita K, Hocart SJ, Coy DH, Jensen RT. Molecular Basis of the Selectivity of Gastrin-Releasing Peptide Receptor for Gastrin-Releasing Peptide. Mol Pharmacol. 2002 Jun 1;61(6):1435–43.

21. Akeson M, Sainz E, Mantey SA, Jensen RT, Battey JF. Identification of four amino acids in the gastrin-releasing peptide receptor that are required for high affinity agonist binding. J Biol Chem. 1997;272(28):17405–9.

22. Ballesteros JA, Weinstein H. Integrated methods for the construction of three-dimensional models and computational probing of structure-function relations in G protein-coupled receptors. Methods Neurosci. 1995 Jan 1;25(C):366–428.

23. Carver JA. The conformation of bombesin in solution as determined by two-dimensional 1H-NMR techniques. Eur J Biochem. 1987 Oct 1;168(1):193–9.

24. Carver JA. A two dimensional 1H NMR study of the solution conformation of gastrin releasing peptide. Biochem Biophys Res Commun. 1988 Jan 29;150(2):552–60.

25. Di Bello C, Gozzini L, Tonellato M, Corradini MG, D’Auria G, Paolillo L, et al. 500 MHz NMR characterization of synthetic bombesin and related peptides in DMSO-d6 by two-dimensional techniques. FEBS Lett. 1988 Sep 12;237(1–2):85– 90.

26. Carver JA, Collins JG. NMR identification of a partial helical conformation for bombesin in solution. Eur J Biochem. 1990;187(3):645–50.

27. Díaz MD, Fioroni M, Burger K, Berger S. Evidence of complete hydrophobic coating of bombesin by trifluoroethanol in aqueous solution: An NMR spectroscopic and molecular dynamics study. Chem - A Eur J. 2002;8(7):1663–9.

28. Erne D, Schwyzer R. Membrane Structure of Bombesin Studied by Infrared Spectroscopy. Prediction of Membrane Interactions of Gastrin-Releasing Peptide, Neuromedin B, and Neuromedin C. Biochemistry. 1987;26(20):6316–9.

29. Cavatorta P, Farruggia G, Masotti L, Sartor G, Szabo AG. Conformational flexibility of the hormonal peptide bombesin and its interaction with lipids. Biochem Biophys Res Commun. 1986 Nov 26;141(1):99–105.

30. Sharma P, Singh P, Bisetty K, Corcho FJ, Perez JJ. Conformational profile of bombesin assessed using different computational protocols. J Mol Graph Model. 2010;29(4):581–90.

31. Valverde A, Gomez-Gutierrez P, Perez JJ. Assessment of the conformational profile of bombesin by computational methods. J Mol Graph Model. 2020 Jul 20;98:107590.

32. Perez JJ, Perez RA, Perez A. Computational Modeling as a Tool to Investigate PPI: From Drug Design to Tissue Engineering. Front Mol Biosci. 2021 May 20;8.

33. Peng S, Zhan Y, Zhang D, Ren L, Chen A, Chen ZF, et al. Structures of human gastrin-releasing peptide receptors bound to antagonist and agonist for cancer and itch therapy. Proc Natl Acad Sci. 2023;120(6):1–12.

34. Isralewitz B, Gao M, Schulten K. Steered molecular dynamics and mechanical functions of proteins. Curr Opin Struct Biol. 2001 Apr 1;11(2):224–30.

35. Sali A, Blundell T. Comparative protein modelling by satisfaction of spatial restraints. J Mol Biol. 1993 Dec 5;234(3):779–815.

36. Shihoya W, Nishizawa T, Yamashita K, Inoue A, Hirata K, Kadji FMN, et al. X-ray structures of endothelin ETB receptor bound to clinical antagonist bosentan and its analog. Nat Struct Mol Biol. 2017 Sep 7;24(9):758–64.

37. Shen M yi, Sali A. Statistical potential for assessment and prediction of protein structures. Protein Sci. 2006 Nov;15(11):2507.

38. Lupala CS, Rasaeifar B, Gomez-Gutierrez P, Perez JJ. Using molecular dynamics for the refinement of atomistic models of GPCRs by homology modeling. J Biomol Struct Dyn. 2018 Jul 4;36(9):2436–48.

39. Cordomí A, Edholm O, Ferez JJ. Effect of different treatments of long-range interactions and sampling conditions in molecular dynamic simulations of rhodopsin embedded in a dipalmitoyl phosphatidylcholine bilayer. J Comput Chem. 2007 Apr 30;28(6):1017–30.

40. Herrera-Hernández MG, Razzaghi N, Fernandez-Gonzalez P, Bosch-Presegué L, Vila-Julià G, Pérez JJ, et al. New insights into the molecular mechanism of rhodopsin retinitis pigmentosa from the biochemical and functional characterization of G90V, Y102H and I307N mutations. Cell Mol Life Sci. 2022 Jan 1;79(1).

41. Schott-Verdugo S, Gohlke H. PACKMOL-Memgen: A Simple-To-Use, Generalized Workflow for Membrane-Protein-Lipid-Bilayer System Building. J Chem Inf Model. 2019;59:2522–8.

42. Case DA, Ben-Shalom IY, Brozell SR, Cerutti DS, Cheatham TE, III, et al. Amber18. University of California, San Francisco; 2018.

43. Maier JA, Martinez C, Kasavajhala K, Wickstrom L, Hauser KE, Simmerling C. ff14SB: Improving the Accuracy of Protein Side Chain and Backbone Parameters from ff99SB. J Chem Theory Comput. 2015 Jul 7;11(8):3696–713.

44. Price DJ, Brooks CL. A modified TIP3P water potential for simulation with Ewald summation. J Chem Phys. 2004 Nov;121(20):10096–103.

45. Vila-Julià G, Granadino-Roldán JM, Perez JJ, Rubio-Martinez J. Molecular Determinants for the Activation/Inhibition of Bak Protein by BH3 Peptides. J Chem Inf Model. 2020;60(3):1632–43.

46. Peralta-Moreno MN, Anton-Muñoz V, Ortega-Alarcon D, Jimenez-Alesanco A, Vega S, Abian O, et al. Autochthonous Peruvian Natural Plants as Potential SARS-CoV-2 Mpro Main Protease Inhibitors. Pharmaceuticals. 2023 Apr 1;16(4):585.

47. Darden T, York D, Pedersen L. Particle mesh Ewald: An N·log(N) method for Ewald sums in large systems. J Chem Phys. 1993;98(12):10089–92.

48. Allen MP, Tildesley DJ. Computer simulation of liquids: Second edition. Comput Simul Liq Second Ed. 2017;1–626.

49. Perez JJ, Tomas MS, Rubio-Martinez J. Assessment of the Sampling Performance of Multiple-Copy Dynamics versus a Unique Trajectory. J Chem Inf Model. 2016;56(10):1950–62.

50. Ryckaert JP, Ciccotti G, Berendsen HJC. Numerical integration of the cartesian equations of motion of a system with constraints: molecular dynamics of n-alkanes. J Comput Phys. 1977;23(3):327–41.

51. Roe DR, Cheatham TE. PTRAJ and CPPTRAJ: Software for Processing and Analysis of Molecular Dynamics Trajectory Data. J Chem Theory Comput. 2013;9:3084–95.

52. Kollman PA, Massova I, Reyes C, Kuhn B, Huo S, Chong L, et al. Calculating structures and free energies of complex molecules: Combining molecular mechanics and continuum models. Acc Chem Res. 2000;33(12):889–97.

53. Tsui V, Case DA. Theory and applications of the Generalized Born solvation model in macromolecular simulations. Biopolymers. 2000;56(4):275–91.

54. Onufriev A, Bashford D, Case DA. Exploring Protein Native States and Large-Scale Conformational Changes with a Modified Generalized Born Model. Proteins Struct Funct Genet. 2004;55(2):383–94.

55. Weiser J, Shenkin PS, Still WC. Approximate solvent-accessible surface areas from tetrahedrally directed neighbor densities. Biopolymers. 1999;50(4):373–80.

56. Gohlke H, Case DA, Biology M, Scripps T, Rd NTP. Converging Free Energy Estimates : MM-PB (GB) SA Studies on the Protein – Protein Complex Ras – Raf. 2003;238–50.

57. Gohlke H, Kiel C, Case DA. Insights into protein-protein binding by binding free energy calculation and free energy decomposition for the Ras-Raf and Ras-RalGDS complexes. J Mol Biol. 2003;330(4):891–913.

58. Shihoya W, Izume T, Inoue A, Yamashita K, Kadji FMN, Hirata K, et al. Crystal structures of human ET B receptor provide mechanistic insight into receptor activation and partial activation. Nat Commun. 2018 Dec 1;9(1).

59. Holst B, Nygaard R, Valentin-Hansen L, Bach A, Engelstoft MS, Petersen PS, et al. A conserved aromatic lock for the tryptophan rotameric switch in TM-VI of seven-transmembrane receptors. J Biol Chem. 2010 Feb 5;285(6):3973–85.

60. Camastra F, Vinciarelli A. Clustering methods. Adv Inf Knowl Process. 2015;131–67.

